# Maps of variability in cell lineage trees

**DOI:** 10.1101/267450

**Authors:** Damien G. Hicks, Terence P. Speed, Mohammed Yassin, Sarah M. Russell

## Abstract

New approaches to lineage tracking allow the study of cell differentiation over many generations of cells during development in multicellular organisms. Understanding the variability observed in these lineage trees requires new statistical methods. Whereas invariant cell lineages, such as that for the nematode *Caenorhabditis elegans*, can be described using a lineage map, defined as the fixed pattern of phenotypes overlaid onto the binary tree structure, the variability of cell lineages from higher organisms makes it impossible to draw a single lineage map. Here, we introduce lineage variability maps which describe the pattern of second-order variation throughout the lineage tree. These maps can be undirected graphs of the partial correlations between every lineal position or directed graphs showing the dynamics of bifurcated patterns in each subtree. By using the symmetry invariance of a binary tree to develop a generalized spectral analysis for cell lineages, we show how to infer these graphical models for lineages of any depth from sample sizes of only a few pedigrees. When tested on pedigrees from *C. elegans* expressing a marker for pharyngeal differentiation potential, the maps recover essential features of the known lineage map. When applied to highly-variable pedigrees monitoring cell size in T lymphocytes, the maps show how most of the phenotype is set by the founder naive T cell. Lineage variability maps thus elevate the concept of the lineage map to the population level, addressing questions about the potency and dynamics of cell lineages and providing a way to quantify the progressive restriction of cell fate with increasing depth in the tree.

**Author summary:** Multicellular organisms develop from a single fertilized egg by sequential cell divisions. The progeny from these divisions adopt different traits that are transmitted and modified through many generations. By tracking how cell traits change with each successive cell division throughout the family, or lineage, tree, it has been possible to understand where and how these modifications are controlled at the single-cell level, thereby addressing questions about, for example, the developmental origin of tissues, the sources of differentiation in immune cells, or the relationship between primary tumors and metastases. Such lineages often show large variability, with apparently identical founder cells giving rise to different patterns of descendants. Fundamental scientific questions, such as about the range of possible cell types a cell can give rise to, are often about this variability. To characterize this variation, and thus understand the lineage at the population level, we introduce lineage variability maps. Using data from worm and mammalian cell lineages we show how these maps provide quantifiable answers to questions about any developing lineage, such as the potency of founder cells and the progressive restriction of cell fate at each stage in the tree.

## Introduction

The cells of developing organisms differentiate into their specialized types by integrating signals from their present surroundings with instructions from their ancestral past. This interplay of mechanisms is reflected in the pattern of phenotypes that emerge in the cell lineage tree [1]. Measurement of this pattern, which involves recording both the phenotypes of, and ancestral relationships between, each cell throughout the lineage tree, results in what is called a lineage map [2]. Lineage maps illustrate the successive bifurcations in phenotypes that underpin a particular differentiation pathway, making them invaluable to experiments investigating the mechanisms involved in fate determination [3]. Development in the nematode *Caenorhabditis elegans* is the classic example of how the lineage map can be used to untangle the roles of pre-programmed instruction and cell-to-cell communication [4–6] in cellular differentiation.

The lineage map allows the common ancestry of cells with shared phenotypes to be identified, thus indicating how deep within the tree a particular cell fate is specified. While fate might not have been specified at a common ancestor itself (lateral inhibition between co-located descendants could be responsible, for example), locating its lineal position is an important step towards finding the mechanisms involved. Interpretation of a lineage map thus starts with identifying the subclones of shared phenotypes. If a phenotype is clonal, meaning exclusive to a subclone, that phenotype can be associated with a single common ancestor; if it is non-clonal, multiple common ancestors were involved (see Table 1). Much of the logic for understanding lineage maps and inferring differentiation pathways from an invariant lineage can be automated [7, 8]. However, in the presence of significant variability, these established techniques become difficult to implement as the procedure of identifying the subclones of shared phenotypes becomes increasingly ambiguous.

**Table 1.** Characteristics of some cell lineage patterns. Organisms are listed in order of increasing complexity. Lineages are characterized in terms of whether cell fate is exclusive to a subclone, the degree of phenotypic variability, and whether there is a way to distinguish between daughters. Lineages from higher organisms are generally unordered, have high variability, and may or may not be clonal.

| *Species* | *Cell origin (tissue)* | *Clonal* | *Variability* | *Ordered* | *Ref.* |
| --- | --- | --- | --- | --- | --- |
| Worm | Embryonic (germ) | ✓ | low | ✓ | [5] |
|  | Embryonic (pharynx) | ✗ | low | ✓ | [5] |
| Leech | Embryonic (epidermis) | ✗ | low | ✓ | [11] |
| Zebrafish | Embryonic (various) | ✗ | high | ✗ | [19] |
| Mouse | Embryonic (various) | ✗ | high | ✗ | [20] |
|  | Lymphoma | ✓ | high | ✗ | [21] |
|  | B-lymphocyte | ✗ | high | ✗ | [14] |

### Variability in cell lineages

The lineage map is a concept born from the study of invariant lineages, such as that for *C. elegans*, where the fixed pattern of phenotypes can, at least in principle, be measured by tracking the progeny of a single founder cell. However, when pedigrees are highly variable, seemingly identical founder cells can give rise to different patterns of descendants. Which of these defines the lineage map? Calculating an average phenotype at each lineal position by pooling across multiple pedigrees can give misleading results since the averaging will suppress the correlations between lineal positions that are so essential for interpreting patterns. Furthermore, the variability between pedigrees, which reflects the potency of founder cells, is an important quantity itself and cannot be represented in a lineage map. While lineage variability is minimal in simple organisms such as *C. elegans* [9, 10] and leech [11], it is greater in higher organisms such as insects and vertebrates [1, 12] and is significant in mammalian cells of clinical importance such as stem cells [13] and lymphocytes [14, 15]. Given the additional variation inherent in molecular-level measurements [16] it is becoming increasingly important to extend the concept of the lineage map to account for variability.

A further problem arises, particularly in higher organisms, when it is not possible to distinguish between the daughter cells from a cell division. This makes the assignment of their relative lineal position arbitrary. Reliably distinguishing between two daughters is possible only when there is symmetry-breaking information available, such as from the orientation of the developing organism. For example, in time-lapse microscopy measurements on *C. elegans*, daughters can be labeled anterior or posterior, dorsal or ventral, left or right depending on their relative positions at the time of division [4, 17]. In higher organisms, however, such symmetry-breaking information often does not exist or cannot be seen. This results in what we will call ‘unordered’ lineages, where there is an ambiguity in the labelling of daughters and, consequently, their subtrees.

A lineage being unordered is not a problem in itself if the phenotype pattern is clear and invariant, since a single complete pedigree measurement represents the lineage map. However, considerable difficulties arise if pedigrees are both variable and unordered. Naive aggregation of multiple pedigrees to get an average phenotype at each lineal position risks suppressing any bifurcation patterns since there is no symmetry-breaking information available to order different pedigrees the same way [18].

Since the majority of pedigree measurements from higher organisms are both variable and unordered [1] (see Table 1), a critical question is whether it is even possible to derive a lineage map from lineage measurements. How do we associate fate specification with fixed lineal positions when the pattern of descendants varies from one apparently identical founder to the next? Clearly a statistical approach is required.

### Previous statistical approaches

A number of statistical methods have been developed to analyze variable, unordered lineage trees. Though these approaches do not directly address the question of how to construct a lineage map, many of them address central aspects of the problem.

A bifurcating autoregressive model [22, 23] was developed to estimate mother-daughter and daughter-daughter correlations using a sample of unordered pedigrees from either *E. coli* or tumor cultures. The model was later used to analyze data from ordered pedigrees to test for lineage asymmetry [24, 25]. This stationary, parametric model allowed for daughters to be conditionally dependent (with respect to their common mother) but forced cousins and more distant relatives to be conditionally independent (with respect to their most recent common ancestor). The subsequent discovery that cousins could be conditionally dependent motivated a theory of cellular inheritance involving chaotic dynamics in lymphoblasts [26]. However, such distant intragenerational dependence might also be interpreted as a delay between fate specification and expression, where a phenotype that has been specified in a mother and its daughters is not expressed until its four granddaughters. These analyses illustrate the importance of having lineages that are large enough, and a model that is general enough, to examine correlations of distant relatives [27]. They also remind us that simple branching process models, which we define to be those assuming conditional independence of daughters, do not properly represent the correlations in a lineage, a fact that was established in early lineage analysis [28, 29]. Although population numbers can be modeled using branching processes [30], allowing for sibling correlations can have important effects on population dynamics [14, 31].

As we indicated earlier, identifying the subtree, or subclone, of shared phenotypes is the first step to inferring where fate is specified. This idea forms the basis of methods to study cell state transitions in bacterial cells or mouse embryonic stem cells [32, 33], where phenotypic similarity among relatives in the same generation was used to infer how much earlier in the pedigree a transition occurred. A similar idea was used in hematopoietic stem cells to measure the multi-generational delay between when an invisible molecular decision occurred and when its effect was expressed as a surface marker [34]. These techniques assume that cell states transition over timescales that are slow compared to the cell cycle duration; alternatively they could be synchronized to cell divisions [35]. Note that, in a lineage map, the generation of a cell is a meaningful quantity, representing the number of divisions since the founder cell, whether that be a zygote, a naive lymphocyte, or some progenitor initiated with a particular stimulus. Thus any model of a developing lineage must be non-stationary.

Several other approaches to statistical lineage analysis have been reported recently. A factor graph method was used to model conditional dependence between daughters [36], with the goal of testing whether pre-programmed instruction or differential cell death was responsible for differentiation of hematopoietic progenitor cells; direct inference of Nanog expression, a pluripotency factor, was used to understand its dynamics in embryonic stem cell lineages [37]; and, a parametric characterization of lineage patterns has been applied to achieve early identification of hematopoietic stem cells [38]. However, these methods are of less relevance to our question of how to build a statistical lineage map.

### Outline

Major efforts are underway to improve the throughput and quality of lineage measurements (see reviews [13, 39–42] and commentary [43, 44]). Recent breakthroughs have resulted in a wealth of data from automated microscopy-based [19, 45–47] and sequencing-based [20, 48–56] techniques. While the technological barriers for these measurements are severe, there are significant barriers to the analysis of the data as well. As we have discussed, there is currently no way to construct a useful lineage map from variable, unordered pedigrees. Since “Central unresolved problems in human biology and medicine are in fact questions about the human cell lineage tree: its structure, dynamics, and variability during development, growth, renewal, aging and disease” [40], generalizing the concept of the lineage map to the population level is of critical importance.

In this paper we provide a solution by proposing lineage variability maps. These involve the variances of, and covariances between, every position in the tree. The supposition is that, to interpret lineage patterns, it is not only the phenotypic values at each lineal position that are important, but also the phenotypic associations between different lineal positions. By developing a generalized spectral analysis for binary trees, we show how to estimate variability maps for a lineage of any depth using measurements from only a few pedigrees. For complete data, our approach is a non-parametric one, involving first and second moments of the data but assuming no distribution function. We could thus, alternatively, refer to these maps as second order lineage maps.

The rest of the paper is organized as follows. Section “Lineage data” describes essential aspects of the data used in this paper. The framework of the model, and how lineal positions are assigned to variables, is given in Section “Analysis framework and labeling conventions”. Section “Covariance estimation”, shows how to estimate all pairwise associations by employing general constraints on symmetry and sparsity. Graphical models are used to create the lineage variability maps and interpret dynamics in Section “Lineage variability maps”. Fate restriction and expression profiles are defined and illustrated in Section “Fate profiles”. A discussion about the interpretations and prospects for this analysis is given in Section “Discussion”.

## Lineage data

Data from 3 types of lineages are analyzed in the paper. Experimental data from T cells provide an example of a lineage with extreme variability and no obvious structure. Previously-published data from *C. elegans* are the example of a lineage with complicated but highly-reproducible structure. Finally, a simulated, stationary branching process provides the benchmark of a featureless, variable lineage and to test the accuracy of the inference procedure. In more detail:

### T cells

New lineage data on CD8^+^ T cells from GFP:OT-1 transgenic mice. Naive cells, expressing a T cell receptor for SIINFEKL peptide from ovalbumin, interact with peptide-pulsed bone marrow-derived dendritic cells to activate clonal expansion [57]. Cells and their descendants are tracked using time-lapse fluorescence microscopy and analysed using custom software [58]. Although multiple phenotypic traits were recorded, in this paper the only trait analyzed is the average area of a dividing cell over its lifetime. Note that only dividing cells were used in the analysis; cells whose fate is unknown, or which died, were counted as missing data. For the early generations used in this study, the numbers of cell deaths were negligible so there was thus no need to account for cell death explicitly. 19 replicate families were used.

### Worm

Published [59] embryonic lineage data from the RW10425 transgenic strain of *C. elegans*. In this strain the *PHA-4* gene for pharyngeal and intestinal tissue is tagged with green fluorescent protein. Gut differentiation occurs early during embryogenesis, with *PHA-4* expression beginning by generations 7 and 8. There are 10 replicate pedigrees.

### Branching Process

Simulated lineages from a stationary branching process. 20 replicate pedigrees are used, with a missing data fraction of 20% assumed. Here we define a branching process to be one where the correlation between mothers and daughters is *h* and daughters are conditionally independent with respect to their common mother. Then, the correlation between any two lineal positions *ς* and *ς* ′ is *h*^*D*^(*ς,ς ′*), where *D*(*ς, ς ′*) is the lineage distance between them. For example, the correlation between sisters is *h* ^2^ and between cousins is *h*^4^ and so on. As will be shown in Section “Lineage variability maps”, the underlying graphical model (of partial correlations) for this branching process is a binary tree. This is generally not the case for real lineages.

Sample lineages from these 3 lineage types are shown in Fig 1 while the expression of each phenotype as a function of generation is shown in Fig 2.

**Fig 1.**
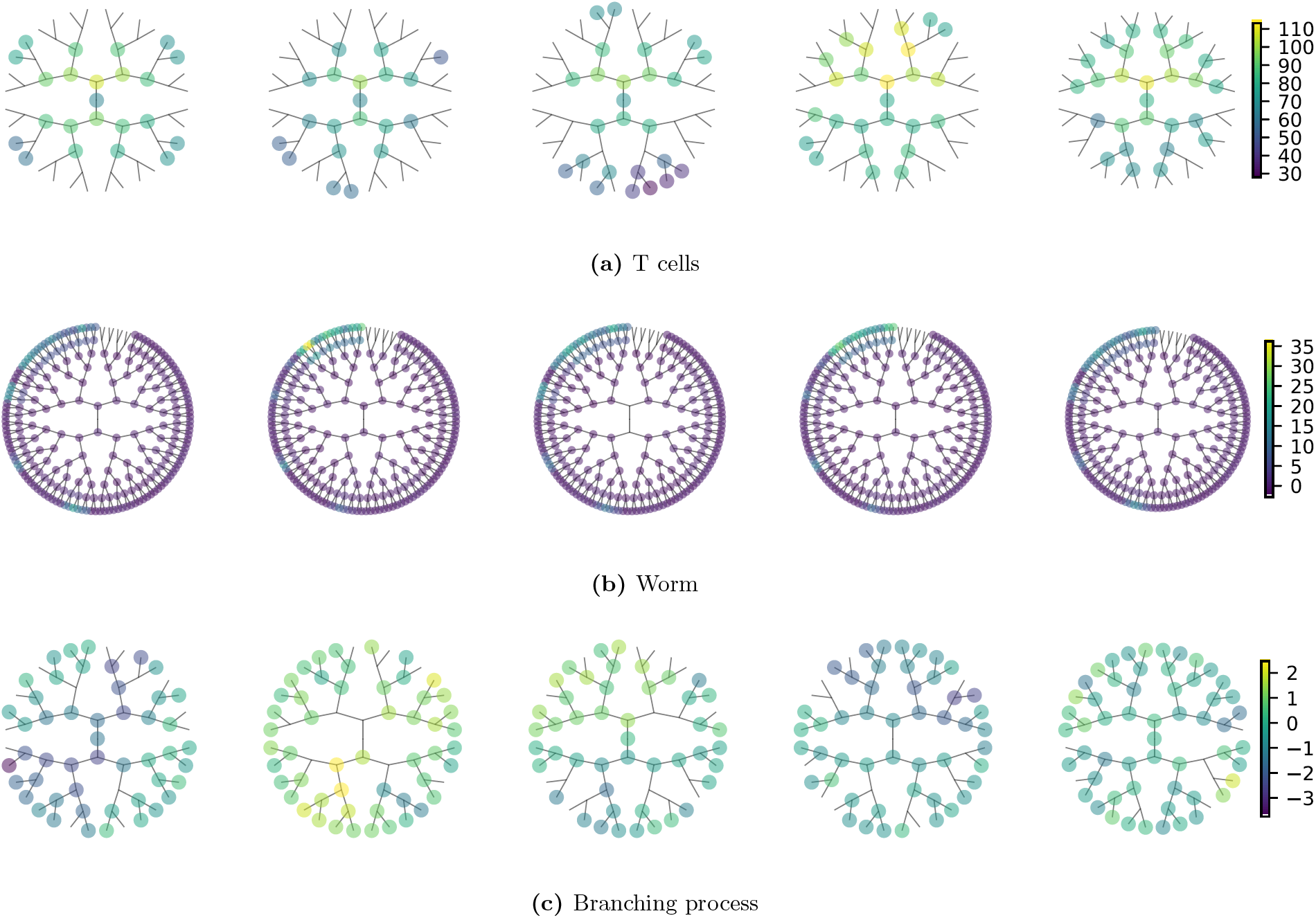
Comparison of some sample lineages. Coloring of the nodes reflects the strength of the trait under analysis (average area over lifetime for T cells, *PHA-4* expression for *C. elegans*). The absence of a node on a branch represents a missing data point. Note that for T cells the root node is the naive cell while for the worm lineage the root node is the zygote (labelled P0 in the *C. elegans* naming convention).

**Fig 2.**
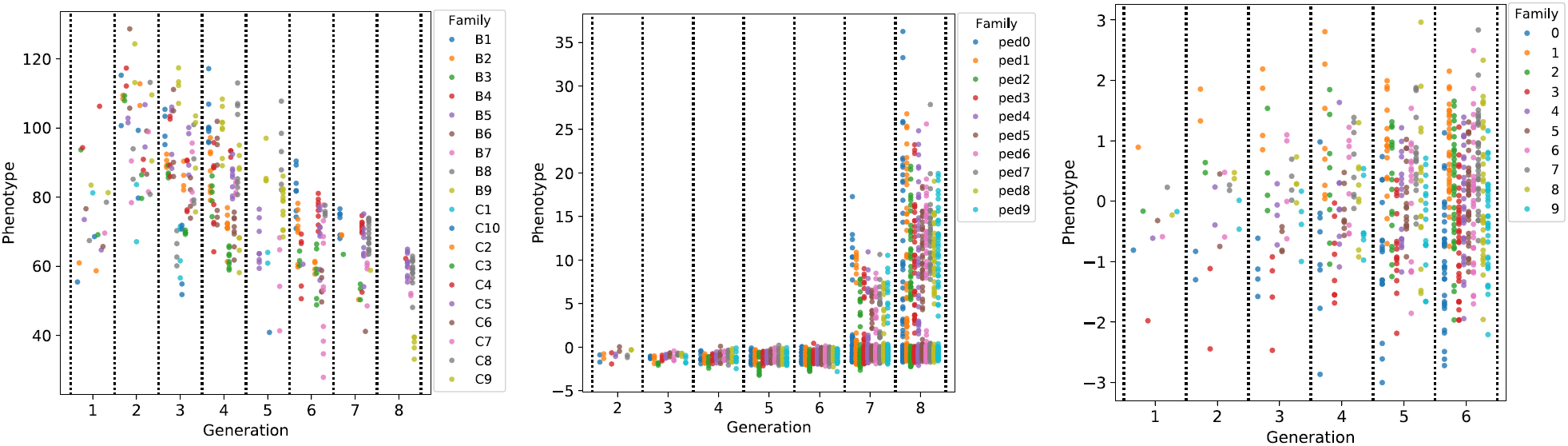
Expression of each phenotype as a function of generation. For T cells the measured phenotype is the average cell area in *µ*m^2^; for *C. elegans* it is the intensity of green fluorescent protein used to tag *PHA-4* expression.

## Analysis framework and labeling conventions

In this study, lineage data are regarded as repeated measurements on pedigrees arising from individual founder cells, each selected at random from a population of similar cells. We restrict our attention to modeling a single trait from pedigrees subject to the same conditions. A sample consisting of multiple replicate pedigrees can then be represented by a two-factor array (*Y*_*ij*_), where *i* has *n* levels corresponding to the number of pedigrees and *j* has *p* levels corresponding to the number of lineal positions within a pedigree. With no meaningful distinctions among pedigrees (they are all of the same cell type and subject to the same conditions) we assume they are independent and identically-distributed replicates. The data can thus be represented by a matrix ***Y*** with *n* replicates (rows) and *p* variables (columns).

Each of the *p* dimensions corresponds to a lineal position. We use a binary number to label each position so that, for example, the first 3 generations are labeled as founder (1), daughters (10, 11), and granddaughters (100, 101, 110, 111), where each label thus encodes the lineal position. We will also need to label generations and subtrees. Generations, *g*, refer to the depth in the tree where we define the founder cell to be at generation *g* = 1. Subtrees are defined by two indices, (*𝓁, τ)*, where *𝓁* refers to the longitudinal coordinate and *τ* to the transverse coordinate of the root node (see Fig 3). By convention, the subtree at *𝓁* = 1 is the entire tree. As we will show, subtrees will be associated with sources of variation. We will need to define a ‘subtree’ (0, 0) that sits outside the lineage and represents variation among lineages. This concept does not exist for a lineage map but is essential for a lineage variability map when different pedigrees may not be the same.

**Fig 3.**
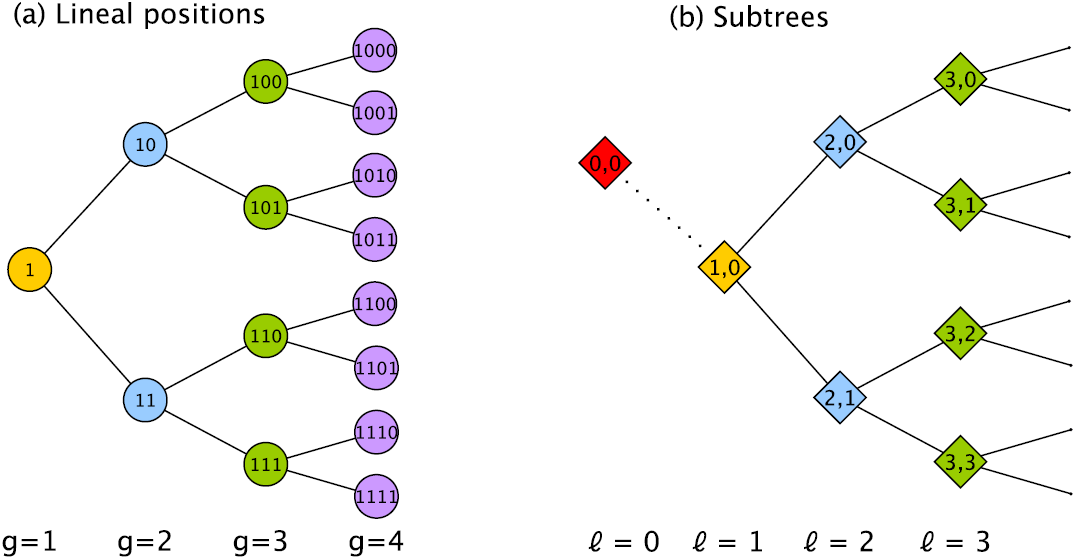
Labeling convention for a lineage tree. (a) Each lineal position is labeled with a binary number. The founder of the tree is located at generation *g* = 1. (b) Each subtree is labeled with two indices (*𝓁, τ)* representing the longitudinal (*𝓁*) and transverse (*τ)* coordinates of its root node. Because, as we discuss later, roots of subtrees are associated with sources of variation we need to create a ‘subtree’ located outside the lineage, called (0, 0), to represent variation among pedigrees. Note that *τ* values are indistinguishable in an unordered tree and will often be ignored.

Often in lineage measurements there are many more lineal positions (*p*) than there are families (*n*). Thus *p ≳ n*, with the disparity getting exponentially worse with the number of generations studied. Performing reliable inference when *p/n >* 1 is an open research question [60]. Best results are achieved when prior knowledge of the problem can be incorporated.

In the next section we describe increasingly more sophisticated steps to reduce the effective dimensionality of the inference calculation, first by exploiting known symmetry properties and then by using observed sparsity properties. Our goal is to identify a scheme where the number of replicates required is independent of the number of generations studied. This is because in practice we might want build maps over many generations from data consisting of only a few pedigrees.

## Covariance estimation

The essential idea for this analysis is to measure second-order variation throughout the lineage by estimating the variance of, and covariance between, every lineal position. This population covariance matrix **Σ** for the lineage involves no assumption about the underlying distribution. It involves just the first and second order moments of the data.

### Unstructured covariance

Let ***y*** be a *p*-dimensional random variable representing the single trait for each lineal position. A naive method for estimating the covariance matrix for ***y*** is to assume it has no structure. This means that only data from the same lineal position in different pedigrees can be pooled. The sample mean 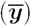 and (biased) sample covariance (***S***) are given by

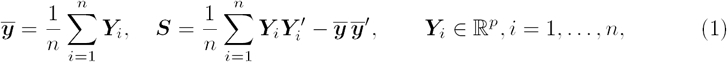

where ***Y***_*i*_ is the data vector from pedigree *i*. This results in the usual estimates of the population mean, ***µ***, and population covariance matrix, 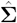,

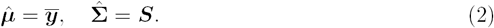

This is not a practical way to estimate **∑** since, as is well known, ***S*** will not be positive definite unless *n* > *p*. To appreciate why this is a prohibitive limitation for lineage data, we examine the complexity of the problem using 3 measures: the effective number of dimensions *p*_eff_, the number of unknown variance-covariance parameters 𝒩_∑_, and the minimum number of replicates *n*_min_ required to ensure 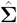 is positive definite. These are given by

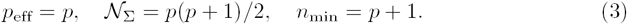

The number of lineal positions for a complete tree of *G* generations is *p* = 2^*G*^ − 1. This means that the number of dimensions, the number of unknowns, and, most importantly, the number of replicates required *n*_min_, increases exponentially with the number of generations being studies. This makes the unstructured covariance matrix impractical for analyzing trees. As we progressively invoke more constraints, we will examine the reduction in these measures of complexity. For example, although *p*_eff_ = *p* for this unstructured case, with group symmetries *p*_eff_ *< p*.

For the analysis to be practical, *n*_min_ should be small and independent of *G*. Then **∑** can be estimated up to any generation *G* with a modest number of pedigrees *n*_min_. To achieve this, our approach is to identify constraints associated with symmetry and sparsity that are specific to the problem of tree-structured variation.

### Symmetry

To understand how symmetry invariance constrains tree-structured variation, we start with intuitive arguments for why certain covariance matrix elements must be equal in an unordered tree. This gives rise to a particular structured form for **∑**. We then describe how the framework of symmetry invariance formalizes this intuition and reveals the independent (orthogonal) components underlying this structured form. The result is a nonparametric spectral analysis for trees that facilitates both inference and interpretation of tree-structured data.

### Structured covariance matrix

To reduce the number of unknowns in **∑**, we begin by identifying a pattern of shared means, variances, and covariances that arise in the unordered tree. This allows pooling of data within a family, in addition to the pooling between families already used in the unstructured covariance estimate.

For the case of first moments, the pattern of shared elements is found by recognizing that, for an unordered tree, some lineal positions are indistinguishable, namely those in the same generation. Equivalently, we could say that the labels identifying members of the same generation are not meaningful. Thus all members within a generation must be assigned the same mean. For example, the mean vector for a 3-generation tree is given by

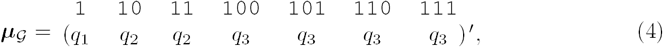

where the subscript 𝒢 identifies the structured mean vector, *q*_*g*_ corresponds to the mean of a cell in generation *g*, and we have explicitly written the cell labels above each element. It is thus apparent that data should be pooled within generations to improve the estimate of these shared means.

Note how, because the tree is unordered, the only information in the first moment of the data is the average of each generation. Other details about the lineage pattern have been lost. *Thus, in unordered trees, we must look at second moments of the data if we want to understand lineage patterns.*

For the case of second moments, the pattern of shared elements is found by recognizing which relationships are indistinguishable. For example, there are two mother-daughter pairs between generations 2 and 3; both must be assigned the same covariance since there is no way to distinguish between the two. We can generalize this intuition by identifying the Most Recent Common Ancestor (MRCA) of a cell pair and adopting a labeling scheme that identifies the generation of each cell and of their MRCA. For example, the pair of cells 10 and 110, which have 1 as their MRCA, should be identified with the 3-index ‘231’, where the first two indices specify the generation of each cell (2 and 3) and the third index specifies the generation of their MRCA (1). Now since the 3-index for another cell pair 11 and 101 is also ‘231’, the two covariances must be equal.

Note how our 3-index scheme identifies the specific generations of both cells and their MRCA, not just the lineage distance between the two cells. This is necessary because, for non-stationary variation in a tree, specific generations are meaningful, not just generational differences. For example, we need to allow for the possibility that sisters in generation 3 have a different statistical association than do sisters in generation 2, even though the lineage distance (between sisters) is the same.

Applying this labeling scheme to each variance and covariance element, the following structured covariance matrix emerges for a 3-generation tree

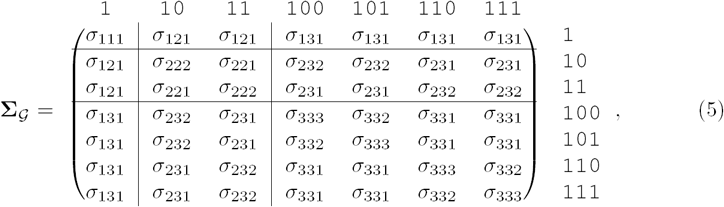

where the subscript 𝒢 denotes a covariance matrix with shared elements. Improved covariance estimation can thus be achieved by pooling across matrix elements with the same 3-index.

Note that the outer product of the structured mean, ***µ***_*𝒢*_***µ***′_*𝒢*_, has a pattern of shared elements that are bounded by the lines in Eq 5. The shared parameters in this less complex pattern are identified by the first two indices of the 3-index in Eq 5. This highlights how **∑**_*𝒢*_ represents the structure of variation that is *in addition to* that due to generational trends seen in Fig 2.

We remark that assuming shared variances and covariances is necessary because, in an unordered tree, we have no information to assume otherwise. We are certainly not assuming that the biology of the lineage tree is symmetric. The need to assume shared parameters for an unordered tree is the same as the need to assume random effects, rather than fixed effects, for batched data when the labels for different batches are not meaningful (see e.g. p.21 [61]).

### Permutation invariance

This pattern of shared means, variances and covariances can be found more formally from symmetry considerations. In general, an object is defined to have a symmetry if it remains invariant under the actions of a group (see Weyl [62] for the classic introduction). A lineage tree has a symmetry because (the action of) permuting daughter subtrees keeps the relationships between lineal positions invariant (see Fig 4). Since no symmetry-breaking information is available (given that the tree is unordered), the permutation has changed nothing about the tree.

**Fig 4.**
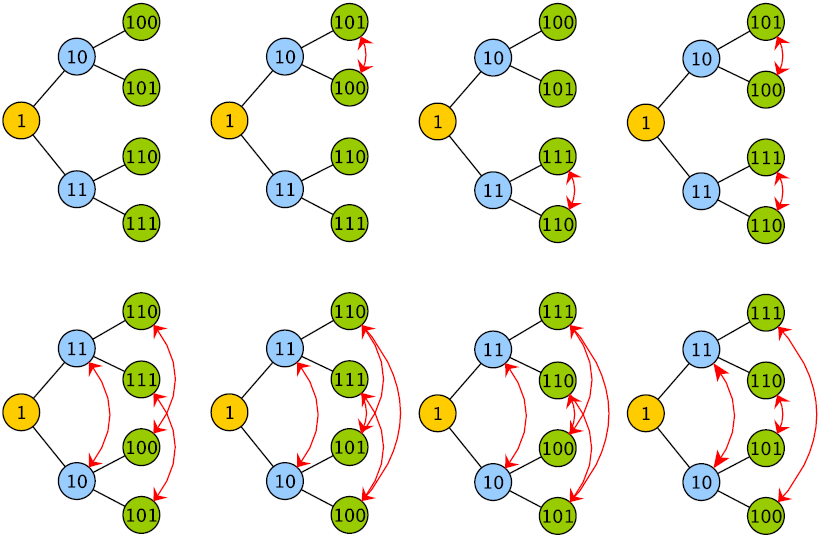
Permutation symmetry of lineal relationships. Certain permutations of lineal positions do not change the relationships in the tree. Here the 8 symmetry-invariant permutations of a tree with 3 generations are shown.

For our purposes, the key idea is that **∑** remains invariant under such permutations of subtrees (since the symmetry group of **∑** is a subgroup of the symmetry group of ***µµ****′* we can focus our attention on the symmetry group of **∑**). Quantifying this intuitive idea involves group representation theory, where matrix multiplications are used to represent symmetry operations [63]. For example, if ***D***_*s*_ is the (*p*-dimensional) permutation matrix representing the action *s* of the group *𝒢*, then the permutation *s* of the variables in ***y*** is represented by ***D***_*s*_***y***. The same permutation of variables in the matrix **∑** is represented by ***D***_*s*_**∑**D′_*s*_, where such conjugation by ***D***_*s*_ is necessary to permute both rows and columns.

The condition that **∑** be invariant under the action of any member of 𝒢 can thus be stated as

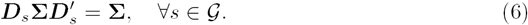

Any symmetry-invariant (i.e. 𝒢-invariant) **∑** thus belongs to the set

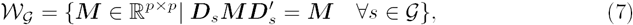

referred to as the fixed point subspace of the group 𝒢 [64]. This is the set of all matrices that are invariant with respect to the group.

### Group-averaged covariance

A standard technique for transforming an unconstrained matrix **∑** into one that is symmetry invariant is the group-average or Reynolds operator (see p. 74 [65]) given by

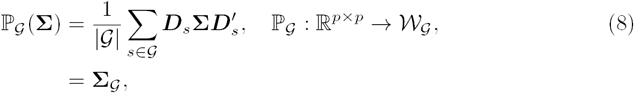

where | 𝒢 | is the order of the group (the number of group elements). This projects the matrix onto the fixed point subspace by averaging over shared elements (referred to as the orbits) of **∑**. It is straightforward to check that the pattern that arises from 𝕡_𝒢_(**∑**), when is the symmetry group of the tree, is the same as that shown in Eq 5. Thus, averaging **∑** over all its allowed permutations (members of the group) generates the properly structured covariance that is invariant to (any further) permutations of the group.

Although this group-averaging approach generates the structured covariance associated with the symmetry group, it is not a practical method for tree-structured data since the number of permutations, | 𝒢 |, grows super-exponentially with *G*. To show this, let *𝒜* be the number of ancestors in the tree, where ancestor refers to any lineal position that has daughters. Let each ancestor be in one of two ‘states’: having its daughter subtrees exchanged or not. For a tree with *G* generations and thus *𝒜 =* 2^*G−*1^ − 1 ancestors, there are 2^*𝒜*^ unique configurations of all ancestor states that keep the lineage relationships invariant. These configurations form the complete set of elements in the group of order 2^*𝒜*^.

Thus, for a 3-generation tree, 𝒜= 3 (corresponding to members 1, 10, and 11) and | 𝒢 | = 2^3^ = 8, where the 8 permutations were shown in Fig 4. For trees with 4 or 5 generations, | 𝒢 | = 128 and | 𝒢 | = 32768, respectively, and the number of permutations quickly becomes unmanageable. Thus the group-averaging approach (Eq 8) is more of a conceptual bridge, connecting the symmetry formalism to the covariance structure, than a practical method for deriving the covariance structure itself.

### Symmetry and generalized spectral analysis

The true benefit of the symmetry formalism is in how it can reduce the original high-dimensional problem into independent lower-dimensional problems that have scientific meaning (see p.161 [66]). This is achieved through a linear transformation from the set of original variables to the set of natural variables defined by the symmetry of the system. The most common example of this is the spectral decomposition of stationary time series data where the underlying symmetry is time invariance and the corresponding natural variables are the Fourier components. Decomposition of a system into its natural variables is thus called generalized spectral analysis, or simply spectral (or harmonic) analysis [66] and has been used in many areas of science and engineering [63].

Formal application of generalized spectral analysis to covariance estimation has been discussed recently [64, 67]. To motivate its application to a complete tree, here we briefly summarize two well-known types of spectral decomposition, Fourier analysis and the analysis of variance (ANOVA), showing how the underlying symmetry of the system defines a linear transformation that diagonalizes the structured covariance matrix.

#### Fourier analysis

Consider 4 variables with the cyclic symmetry shown in Fig 5a. These could be, for example, variables in a temporal sequence where the absolute value of time is not meaningful. Such time invariance means that the covariance matrix does not change if the variables are cyclically shifted, as long as there is no change in their relative ordering. Variation in this set of variables is regarded as stationary since only the *differences* between variables matter, not their absolute position. The covariance matrix then has a circulant structure

**Fig 5.**
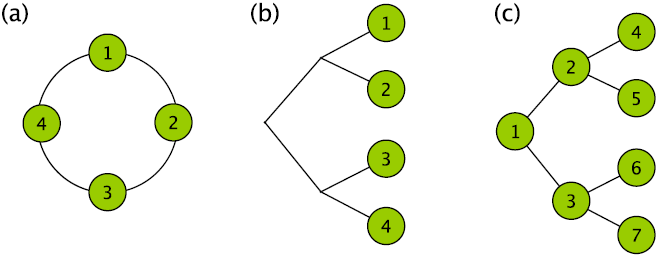
Cyclic and tree-structured symmetries. (a) A cyclic symmetry structure is one that remains invariant under a shift of all the variables (around the circle in the figure shown) that preserves their relative ordering. This cyclic symmetry defines the discrete Fourier transform. (b) A tree symmetry structure is one that remains invariant under permutations within groups and permutations of groups. This symmetry gives rise to the analysis of variance for nested pairs and also defines the Haar wavelet transform. It is applicable when it is just the leaf nodes that are of interest. (c) When all the nodes of a tree are of interest, the underlying symmetry is still that for the tree. The associated transformation is derived in this paper and discussed in the next section.

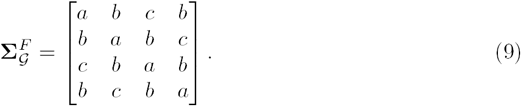

It is well-known that the circulant structure defines a unitary transformation matrix called the discrete Fourier transform (DFT) matrix which, for 4 variables, is given by

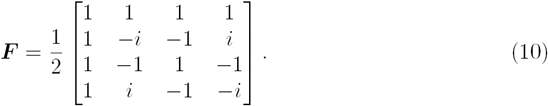

Each column represents a natural variable of the cyclic symmetry, better known as a Fourier basis vector. Using ***F*** to transform 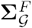 into this natural basis results in a diagonal matrix called the spectral covariance, where the diagonal elements represent the spectrum. Thus, the circulant-structured matrix is transformed into the spectral covariance using the DFT matrix.

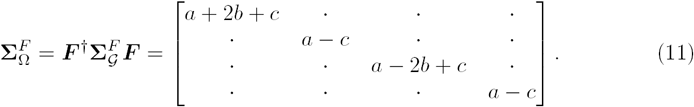

#### ANOVA on nested pairs (Haar wavelet analysis)

Now consider the problem of nested batches of variables, a standard problem in the analysis of variance, or variance components analysis. Consider the case of 2 batches each containing 2 variables. This can be depicted as leaves on a binary tree as shown in Fig 5b. The symmetry operations for this structure are the permutations within groups and permutations of groups, or, as we discussed earlier, the exchange of daughter subtrees. The covariance matrix invariant under these symmetry operations has the form

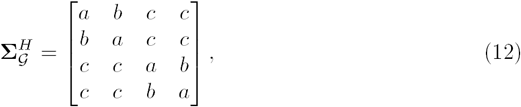

which was given in the bottom right corner of Eq 5. The matrix that diagonalizes this structure,

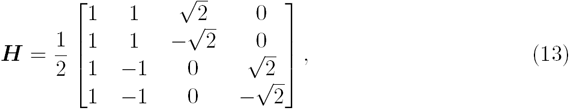

is known as the Haar (wavelet) transform matrix. Each column defines a natural variable of the tree symmetry and represents a source of variation or a wavelet component. Using ***H*** to transform 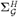 into this natural basis results in a (diagonalized) spectral covariance

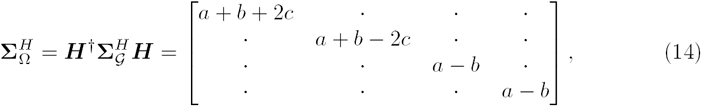

where the diagonal elements are known as the components of variance (if we regard this from the ANOVA perspective), or the Haar wavelet spectrum (if we regard this as wavelet analysis). Here there are 3 sources of variation: between trees (*a* + *b* + 2*c*), within trees (*a* + *b −*2*c*), and within subtrees (*a−b*).

We emphasize that the change-of-basis matrices ***F*** and ***H*** are defined by the symmetry of each system. They transform the original variables into a set of non-interacting natural variables (Fourier or Haar wavelet components) which define the meaningful components of variance. It was Tukey [68] who first showed that Fourier decomposition can be regarded as a branch of variance components analysis.

It is worth pointing out how this diagonalization, or eigendecomposition, of the covariance matrix, relates to traditional principal components analysis. In generalized spectral analysis, the eigenvectors (given by the columns in ***F*** and ***H***), or, more precisely, the eigenspaces, are determined by the *structure* of **∑** and do not depend on its entries. In addition, the eigenvalues are linear functions of the entries. Neither of these properties are true, in general, for principal components analysis.

### Generalized spectral analysis of a complete tree

Having examined the case of a tree where only the leaf nodes are of interest (Fig 5b), we now examine the case where all positions in the tree are of interest (Fig 5c). For the complete tree, we already know the structured covariance **∑**_*𝒢*_ (see Eq 5). Our tasks then are to derive the change-of-basis matrix, interpret the natural variables, and calculate the spectral covariance.

The derivation of the change-of-basis matrix, ***T***, for a complete tree is shown in Appendix A2. This represents the generalization of the Haar transform matrix ***H*** to a complete tree. For a 3-generation tree it is given by

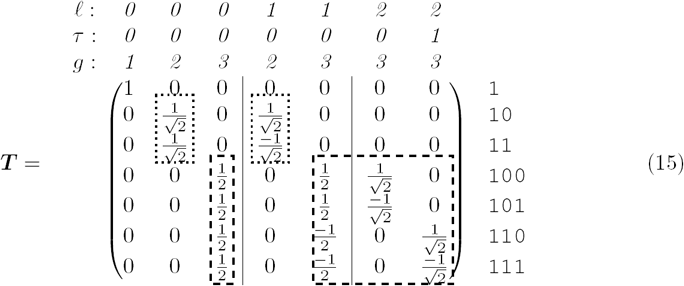

where the columns, as usual, define the natural variables. There are two equivalent ways of interpreting these natural variables: from the ANOVA perspective, and from the wavelet perspective. It is useful to consider both.

From the nested ANOVA perspective, each natural variable is associated with a source of variation (*𝓁, τ,)* located at the root of a subtree. Because we are considering more than one generation, we must also specify the generation *g* in which the variation is observed (see Fig 3 for the labeling convention). From the wavelet perspective, *𝓁* represents the transverse scale of the variation, *τ* the transverse position, and *g* is the generation in which the variation is observed. The 3-index label for each natural variable is given above each column in Eq 15, with vertical lines used to partition the different *𝓁*.

The change-of-basis matrix is straightforward to extend. For example, a tree with 4 generations gives

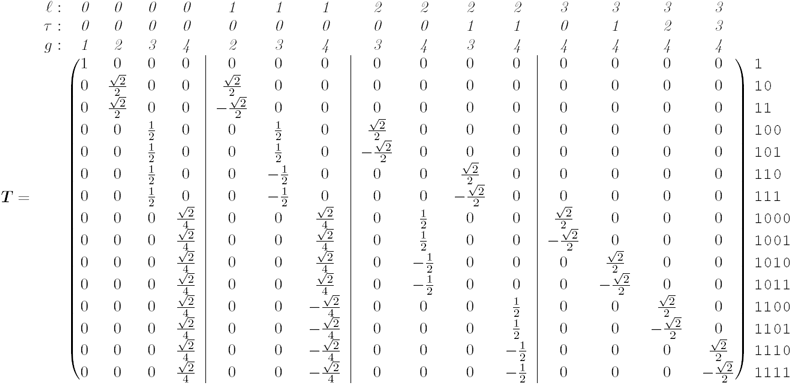

A visual representation of how these natural variables are constructed from the original variables is shown in Fig 6 for the case of a 4-generation tree. This emphasizes how the *𝓁*-coordinate of the source of variation characterizes the scale of the pattern. Fig 7 shows a few examples of the natural variables to illustrate how they are convenient, elemental components for describing tree-structured variation.

**Fig 6.**
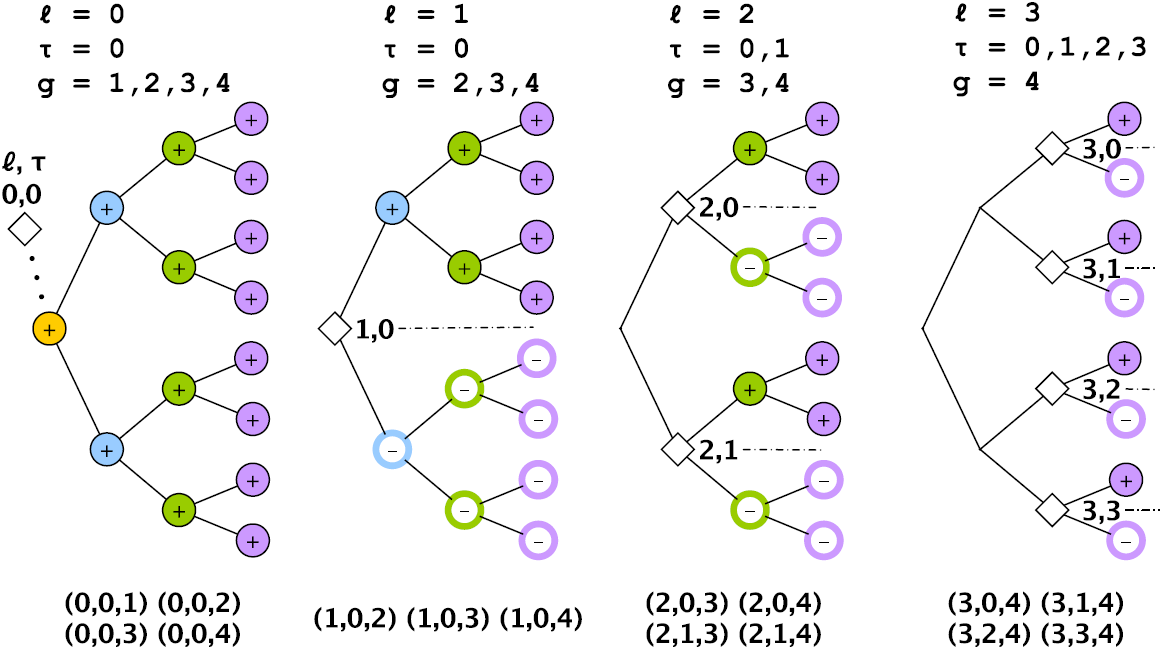
Construction of the natural variables for a tree with 4 generations. Each natural variable is identified by a source of variation (*𝓁, τ)*, corresponding to the root of a subtree, and a generation *g*. The + *−* and at each lineal position illustrate how the original variables are combined to form a natural variable. The 15 natural variables thus defined by the 3-tuple (*𝓁, τ, g*) are listed in the bottom row. Since the *τ* coordinates are indistinguishable, only 10 of the natural variables (those with *τ* = 0, say) are unique.

**Fig 7.**
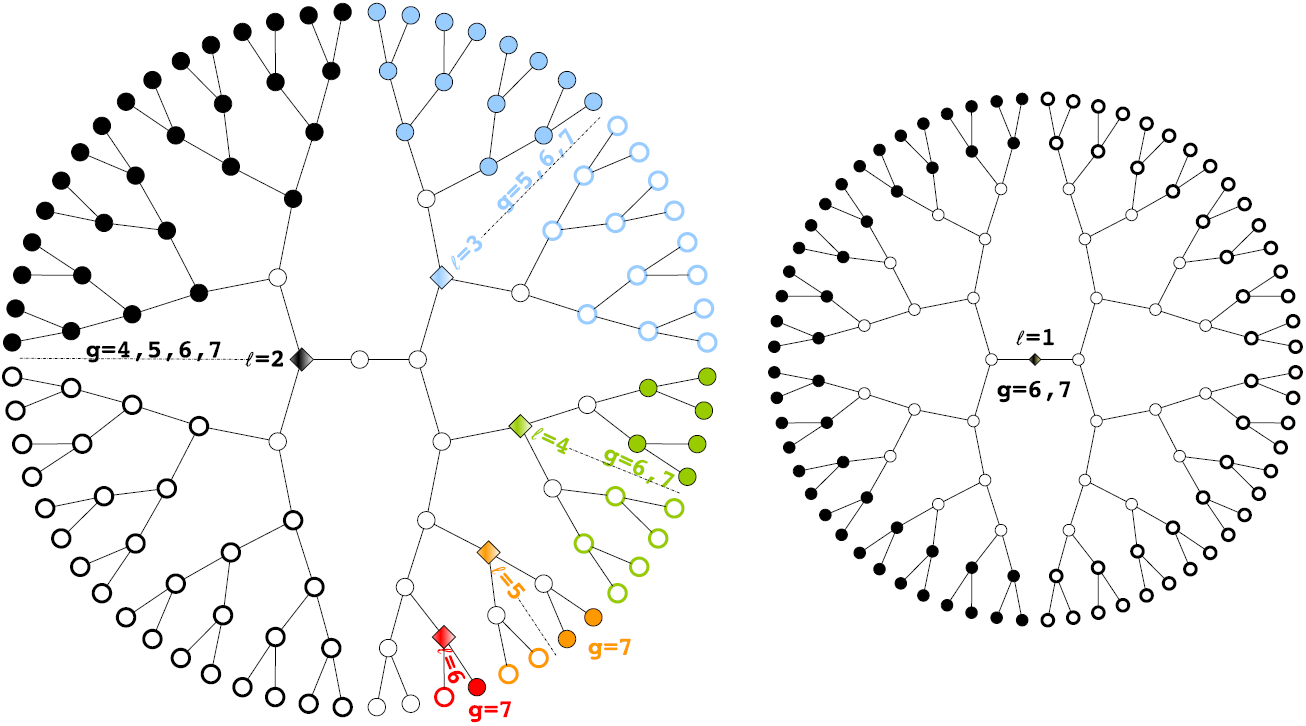
Bifurcated subtrees. Patterns on a tree can be described in terms of natural variables, or elemental components, examples of which are shown here. Each component is a bifurcated pattern centered on a subtree (*𝓁, τ)* and expressed in a generation *g* (where *τ* is ignored in an unordered tree). For example, the blue/non-blue bifurcated pattern is centered on subtree *𝓁.*= 3 and observed at generations 5, 6, and 7. Note that *𝓁* = 1 variation (on the right) is a bifurcation across the whole pedigree, while *𝓁.* = 0 (not shown) represents variation among different pedigrees.

The natural variables are not particularly surprising: they are just those one would define in a nested ANOVA or Haar wavelet analysis if each generation were considered separately. Perhaps more surprising is their arrangement in ***T*** : although Eq 15 contains every column of the Haar transform matrix for generations 2 (dotted lines) and 3 (dashed lines), these matrices are not incorporated simply as a direct sum. Instead, representation theory demands that we group the natural variables by (*𝓁, τ)*. When we do this and apply ***T*** to **∑_*G*_** from Eq 5 we get a *block*-diagonalized spectral covariance,

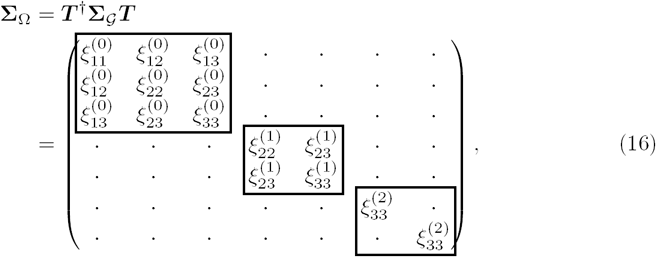

where each block corresponds to a source of variation *𝓁* and its associated generations *g*, where *g > 𝓁*. Here we label matrix elements as 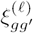 where subscripts refer to the pair of interacting generations, *g* and *g′* (there is no need to use *τ* as a label since elements differing only in *τ* have identical values). Note how the components of variation that we encountered on the diagonal in 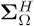 (Eq 14), where only third generation variables were of interest, are here labeled as 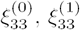, and 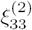 They are still on the diagonal but are grouped with their counterparts from generations 1 and 2.

To better appreciate the block-diagonal structure of **∑**_Ω_, we show it as a heat map for the case of a 4-generation tree (Fig 8b) along with the corresponding **∑**_*𝒢*_ (Fig 8a). This emphasizes how each block *𝓁* is further block-diagonalized by *τ.* In the terminology of group representation theory, *𝓁* identifies an isotypic subspace while *τ* identifies an irreducible subspace - a subset of the isotypic subspace. In Fig 8b, the isotypic blocks are bounded by dashed lines, while the irreducible blocks are bounded by dotted lines.

**Fig 8.**
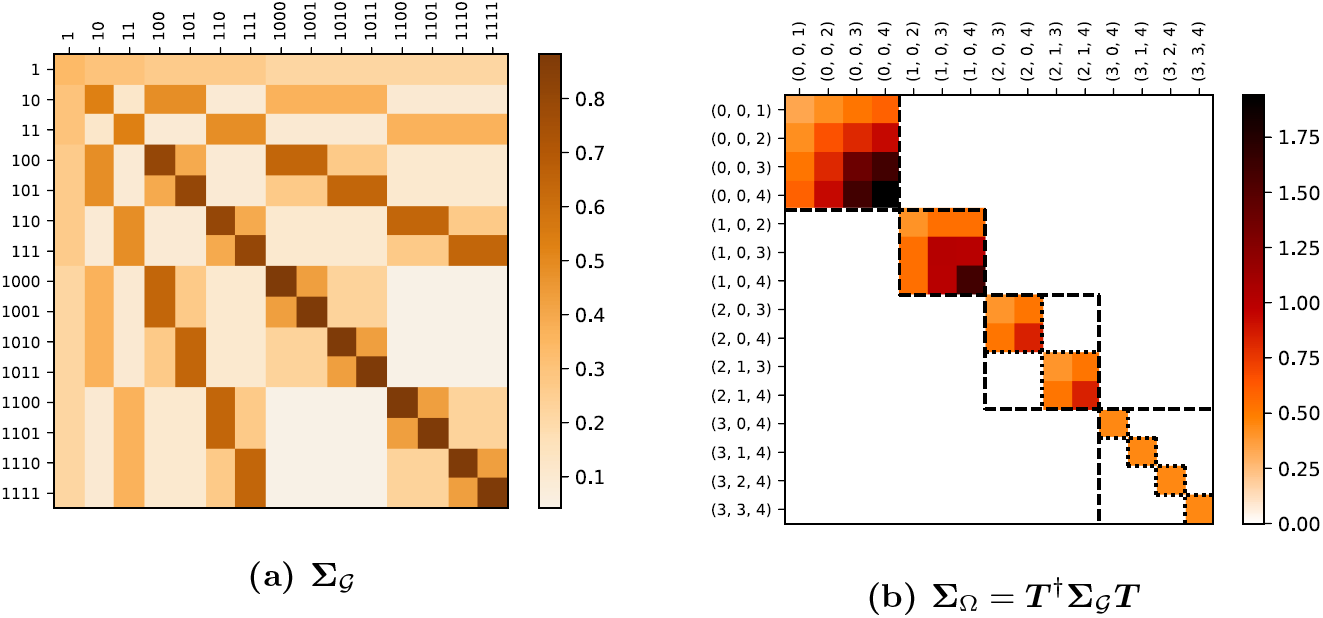
Heat maps of ∑_*𝒢*_ and ∑_Ω_ for a complete tree. This example was taken from the first 4 generations of the branching process. Natural variables along the axes of **∑**_Ω_ are given in the format (*𝓁, τ, g*). Isotypic blocks are bounded by dashed squares and correspond to a given *𝓁*. Irreducible blocks correspond to a source of variation (*𝓁, τ)* and are bounded by a dotted square. For *𝓁* = 0 and 1 the isotypic and irreducible blocks coincide since there is only one *τ* index.

The primary benefit of identifying the spectral transformation for the complete tree is that **∑**_Ω_ contains all the information in **∑**_*𝒢*_ but in a much simpler form. Having pooled the data to obtain 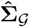 G one simply performs the linear transformation to get 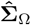.

We pause briefly to examine how this generalized spectral analysis for a complete tree is analogous to traditional Fourier analysis for a time series. As we mentioned, bifurcated subtrees are the natural variables for a binary tree and are thus analogous to sine and cosine waves. Any pattern on a tree, whether or not it is clonal, can thus be defined as a superposition of bifurcated subtrees. This idea is useful when trying to interpret non-clonal lineage patterns: whereas a clonal pattern is associated with a single subtree, a non-clonal pattern is a superposition of multiple subtrees.

Another analogy is between the ordering of the tree and the phase of a time series. Our ability to average different trees regardless of their ordering is similar to the ability to average the spectra of different time series having unknown starting phases. Here one knows that to detect structure in the time series, one should average their spectra, not the time series themselves. Other analogies are shown in Table 2.

**Table 2.** Generalized spectral analysis. Well-known quantities in Fourier analysis have their direct analogs in the spectral analysis of a tree.

| *Fourier analysis* | *Tree analysis* |
| --- | --- |
| Sine, Cosine waves | Bifurcated subtrees |
| Phase | Ordering of the tree |
| Auto-covariance | Structured covariance, $\Sigma_G$ (Eq 5) |
| Discrete Fourier Transform matrix | Change-of-basis matrix, $T$ (Eq 15) |
| Power spectrum | Spectral covariance, $\Sigma_\Omega$ (Eq 16) |

### Complexity of the structured covariance

Spectral decomposition shows that the high-dimensional covariance estimation problem involving shared parameters in 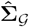 G is equivalent to several, lower-dimensional covariance estimation problems given by the irreducible blocks in 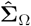 We can use this to calculate the complexity of 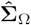 as we did for the unstructured covariance (Eq 3).

Because each unique irreducible block is an independent, unstructured estimate of a covariance matrix, the effective number of dimensions, *p*_eff_, is given by summing the number of dimensions for each *unique* irreducible subspace. The number of free parameters in the covariance matrix, *N* _∑_, is found by summing the number of parameters in each *unique* irreducible block. The minimum number of replicates required, *n*_min_, is found from the dimensionality of the largest irreducible block (*𝓁* = 0). Thus

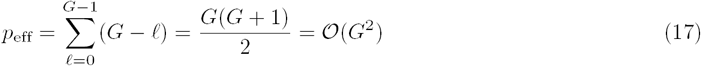

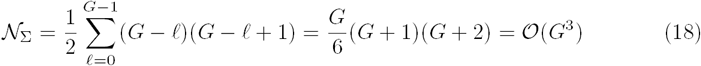

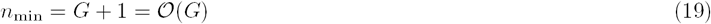

The group-symmetric model is thus significantly more constrained than the unstructured model, with the number of parameters growing polynomially with *G* instead of exponentially (compare Eq 3). Note how *p*_eff_ *< p* (when *G* ≥ 3), a reduction in the effective number of dimensions that was not apparent from **∑**_*𝒢*_ alone.

Nevertheless, even with these symmetry constraints, *n*_min_ still grows with *G*, albeit linearly (Eq 19) instead of exponentially (Eq 3). This means that, for a fixed set of *n* replicates, there will always be a limit to the number of generations that can be analyzed. We need an additional constraint.

### Sparsity

The additional constraint comes from recognizing that the *G − 𝓁* natural variables in each irreducible subspace (*𝓁, τ)* represent a time series from generation *𝓁.*+ 1 to *G* (see Section “Generalized spectral analysis of a complete tree”). Together, the unique irreducible subspaces comprise a set of *G* independent time series each starting at a different generation but all ending at *G*. A standard technique for imposing structure on a time series is to consider it a fixed order Markov chain.

Before doing this, we first need to justify some properties of the inverse covariance, or precision, matrix ***K*** = **∑**^*-*1^. In particular, because **∑**_*𝒢*_ is 𝒢-invariant, its inverse ***K***_*𝒢*_, has the same structure [69]. This means that the spectral precision matrix, ***K***_Ω_ = ***T*** ^*†*^***K***_*G*_ ***T*** has the same block-diagonal structure as **∑**_Ω_. Hence each irreducible block 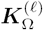 in the spectral precision matrix is just the inverse of the corresponding irreducible block in the spectral covariance 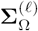:

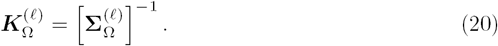

The problem of imposing a Markov constraint on 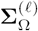 is thus one of imposing sparsity on 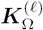. More specifically, matrix elements in 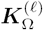 outside a diagonal band (the tri-diagonal in the case of a 1st order Markov process) are constrained to be zero. Remember that it is the structure of each 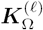 that is sparse; the precision matrix itself, ***K***, may not be particularly sparse. We remark that a zero in the precision matrix enforces conditional uncorrelatedness between two variables without assuming Gaussianity (if the distribution is Gaussian, then this pair of variables is also conditionally independent).

A restricted-order Markov chain is a simple case of a decomposable graphical model [70, 71] and thus yields an explicit estimate of the covariance matrix. Following the procedure for a decomposable model, we organize variables in the irreducible block into cliques and separators, a straightforward exercise for a Markov chain of any order. If 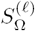 is the (unstructured) estimate of the irreducible block, we label sub-blocks of cliques and separators within 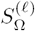 as

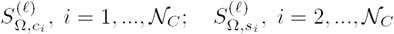

where the subscript *c*_*i*_ refers to a clique, *s*_*i*_ refers to a separator, and *N* _*C*_ is the number of cliques in the irreducible block. The covariance estimate for an irreducible block is then given by (p.145 [71])

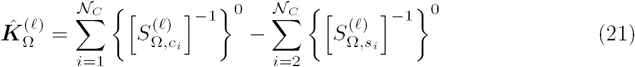

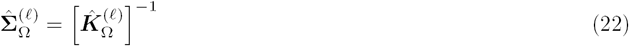

where the expression {*ϒ*}^0^ denotes a matrix with the dimensions of 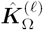 which has its appropriate sub-block occupied by *ϒ* and zeros elsewhere.

This expression makes it clear that, since it is the inverse of the clique and separator sub-blocks that are required, it is only these sub-blocks (with maximum dimension *ℳ* + 1) that need to be positive definite. The minimum number of replicates required for positive definiteness is thus set by the order *ℳ* of the Markov process, which is fixed, rather than by the size of the irreducible block, which grows linearly with *G*. In general then, *n*_min_ = *ℳ* + 2 and we have finally achieved our goal of having the data requirements be independent of the number of generations being analyzed. Note that restricting the non-zero parameters in the precision matrix to be on the diagonal band means that *N* _∑_ ∼ 𝒪 (*G*^2^), down from the cubic dependence in Eq 18. *p*_eff_ remains unchanged.

Inspection of the T-cell and worm lineage data show that, at least up to generation 4, non-zero values in 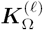 are indeed primarily confined to the tri-diagonal. This justifies the (first-order) Markov process assumption, and we hereafter use it to extend the analysis to higher generations.

### Missing data

The covariance estimates described above assume complete data. In reality, some measurements are missing, often because data collection is imperfect but also because cells die and have no descendants (although in the datasets analyzed in the paper, cell death is essentially negligible).

A simple solution is to apply the Expectation-Maximization (EM) algorithm [72], assuming a multivariate Gaussian to impute the missing data. Before describing how we do this, we remark that the covariance estimation procedure we have described thus far is distribution-free, providing a non-parametric estimate of second-order variation. It is only to account for missing data that we invoke a distributional assumption. In Appendix A3 we show that the maximum likelihood estimate (MLE) for a multivariate Gaussian with the symmetry and Markovian constraints discussed above is in fact the covariance estimate we have already found. It is thus straightforward to apply the EM algorithm with a multivariate Gaussian to address the missing data problem.

The EM algorithm iteratively improves the estimate of the covariance matrix, generating expected values of the sufficient statistics at each step. In the E-step, the current estimate of the mean 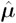 and covariance matrix 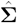 are used to calculate the expected sufficient statistics for each replicate, conditioned on the observed data. The average *Š* over all replicates is then calculated. In the M-step, *Š* is used in the MLE calculation of the irreducible blocks (as described above) to update the estimate 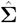. The E and M steps are then repeated until 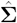 converges.

In more detail (p.223 [73]), the first and second order statistics are calculated for each replicate *i* by partitioning the variables into observed sets, labelled *o*_*i*_, and unobserved sets, labelled *u*_*i*_. Members of each set usually differ from one replicate to the next. The vector of unobserved values in each replicate is then filled by its expected value conditioned on the vector of observed values:

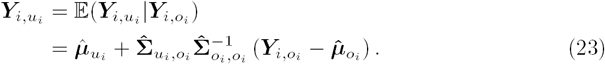

Combining these with the observed values completes the first order statistic, 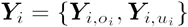 for *i*.

The second order statistic (***Y Y)***_*i*_ for each replicate *i*, partitioned into observed and unobserved sections, is found from

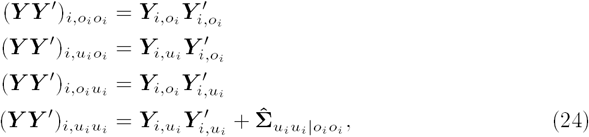

where

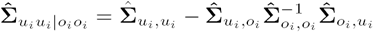

is the residual covariance of the unobserved variables after conditioning on the observed variables.

Once this exercise has been completed for all replicates, the sample mean and covariance are calculated from the usual

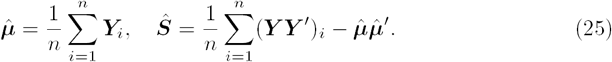

The estimated sample covariance, *Ŝ*, is then used in the procedures described in the previous sections to calculate a new estimate, 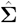 Iterating these steps gives the following algorithm:

1. Initialize 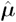 and 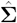.
2. Expectation step to determine the expected value of the sufficient statistics for each replicate. Use Eqs. 23, 24, 25 to calculate the updated estimate 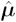 and the estimated sample covariance, *Ŝ*
3. Maximization step to find 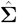 from *Ŝ*
  a. Find 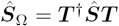.
  b. Set elements outside the diagonal blocks to zero.
  c. If there is more than one irreducible block in a given isotypic block, average them and assign the result to all of them.
  d. For each unique irreducible block, find 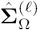 from 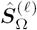 using Eq 21 and 22, assuming a Markov chain of given order *ℳ*.
  e. Recover 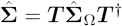
4. Return to Step 2 until convergence.

Note that here, rather than pooling matrix elements in **Ŝ** to estimate **Ŝ** _𝒢_ and then spectrally transforming the result to get the block-diagonalized *S*_Ω_, we instead spectrally transform *S* and perform the averaging the the spectral domain (steps 3b and 3c) to get *S*_Ω_. The two approaches give identical results.

#### Lineage variability maps

Our focus thus far has been to estimate ∑ for the complete tree. The approach we described can in principle be applied to lineages with any number of generations and needs only a few replicates (pedigrees) to ensure positive definiteness. For the rest of the paper we turn to the problem of interpreting ∑.

In this section we visualize 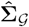 and 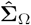 using graphical models to produce different ‘maps’ of the variation in the lineage. We call these lineage variability maps. For 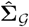 we use undirected graphs, since lineal positions within a generation have no ordering, and we call the result a lineage correlation map.

For 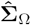 we can use directed graphs, since natural variables belonging to an irreducible block are ordered in a sequence. Thus the spectral transformation allows the undirected graph to be converted into a directed one. This graph, which we call a dynamic lineage map, compactly represents the dynamics of the bifurcated expression pattern in each subtree.

#### Lineage correlation map

To visualize the network of statistical associations between different lineal positions we use undirected graphs [70, 71] defined either by marginal or by conditional associations. For the network of marginal associations the strength of an edge between a pair of variables is defined by the Pearson correlation coefficient, 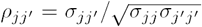 where σ _*jj*_ ′ is an element of 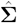. For the network of conditional associations the strength of an edge is determined by the partial correlation 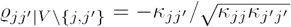 where *κ*_*jj*_′ is an element of 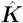, and *V* {*j, j′*} refers to the set of variables excluding *j* and *j′*.

Both types of undirected graphs are shown in Fig 9 for the 3 lineage types. The network of conditional associations identifies direct interactions between variables, conditioned on all other variables, and, as expected, generally provides a sparser representation than does the network of marginal associations.

**Fig 9.**
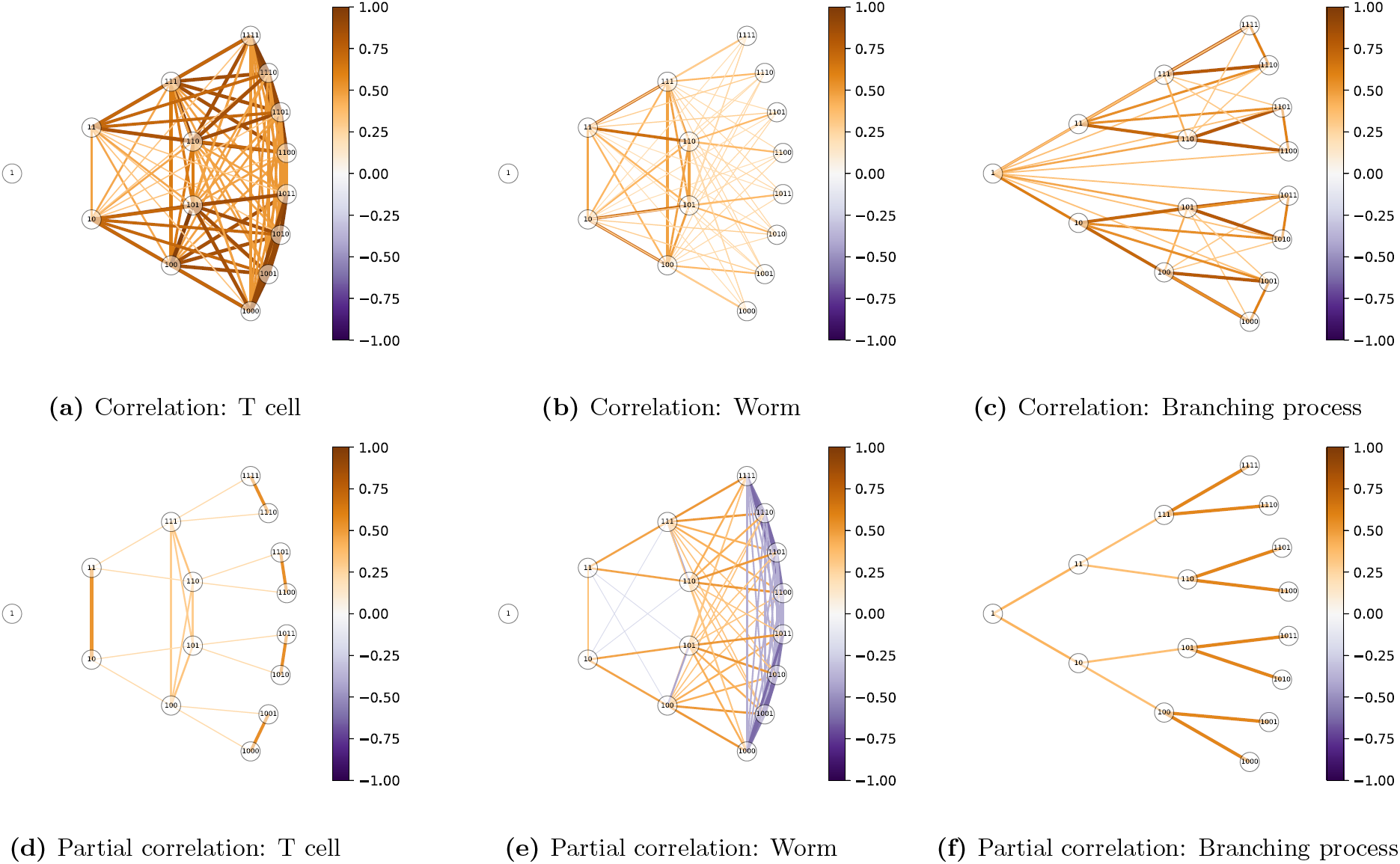
Lineage correlation maps. These are undirected graphs in the original variables. The color of edges in each graph corresponds to the correlation (top row) or partial correlation (bottom row) between pairs of lineal positions. To avoid clutter only the first 4 generations are shown. Note how the graph of partial correlations (9f) for the simulated branching process, where daughters are conditionally uncorrelated, is a binary tree. This is not the case for the real lineages.

Note how a binary tree is revealed in the graph of partial correlations for the branching process (Fig 9f). This is expected since our branching process defined daughters to be conditionally uncorrelated. In the network of partial correlations this assumption reveals itself as the lack of an edge between sisters. In contrast, in the partial correlation graphs for T-cell (Fig 9d) and worm (Fig 9e) lineages, sisters are often joined by edges. This arises when the correlation between sisters is greater or less than the squared correlation between mother and daughter, a long-documented observation in cell lineages (see e.g. [28, 29]). This is the simplest demonstration of the fact that phenotypic variation in real lineages cannot be modeled as a branching process.

The graphs in Fig 9 allow us to examine how the network of *phenotypic* associations compares with the network of *lineal* relationships; though the latter is a binary tree, the former may not be. This emphasizes that, although we must assume that phenotypic variation in an unordered tree has the *symmetry* of a binary tree, we do not assume it has the *sparsity* of a binary tree.

### Dynamic lineage map

A problem with representing each lineal position as a node is that the graph appears cluttered since there are many edges and nodes with similar strengths. This problem gets exponentially worse with increasing generations. Such redundancies disappear when examining the tree over its natural variables.

Since the natural variables in each irreducible subspace are ordered by generation they can be represented by a directed graph [74–76], with each variable conditioned on the past. Each irreducible subspace is thus a chain representing a subtree *𝓁*, with the complete tree thus being represented by *G* independent chains. In the language of graph theory, the tree is composed of connected components, each of which is a chain. Each chain describes how the bifurcated expression pattern associated with a subtree *𝓁* propagates through subsequent generations.

The structural equation, or causal, model underlying each chain is a non-stationary time series given by the following system of equations:

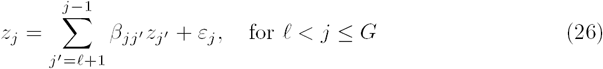

Note that each irreducible subspace is represented by its own system of equations but we avoid the superscripts *𝓁* to reduce index clutter. Here *z*_*j*_ is a natural variable corresponding to a generation *j, β*_*jj*_′ is the regression coefficient of generation *j* on *j*^′^, and *ε*_*j*_ is an independent random variable with a mean of zero representing variation originating at generation *j* that has expected variance 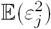. Defining a lower-triangular coefficient matrix ***B*** = (*b*_*jj*_′) gives the system of equations in matrix form:

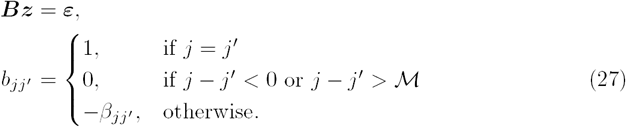

The structural equation model parameters *β*_*jj*_′ and 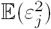 can be found using a modified Cholesky decomposition of each 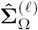,

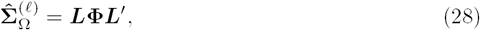

where **Φ** = (*ϕ*_*jj*_′) is diagonal and ***L*** is lower triangular. Then since 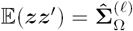, we find that ***L***^*-*1^ = (*b*_*jj*_′). This means that *β*_*jj*_ ′ can be found using Eq 27 and 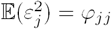.

The directed graph can then be defined with edge weights given by *β*_*jj*_*′* and node strengths given by 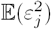. The edges represent transmission of variation while the nodes represent innovations. If *|β*_*jj*_′ *| <* 1 then transmission is regressive, with descendants gradually losing memory of previous generations. However, if |*β*_*jj*_′*| >* 1 then variation from source (*𝓁*) observed at generation *j*′ is *amplified* during transmission to generation *j*. Thus large variation can either arise directly from a large innovation or it can be the result of strong amplification of small variation (or both).

These directed graphs compactly summarize the dynamics of phenotypic variation throughout the lineage. Examples for the 3 lineages types are shown in Fig 10. Each connected component, given by a row, represents how the bifurcated expression pattern associated with a subtree *𝓁* propagates down successive generations.

**Fig 10.**
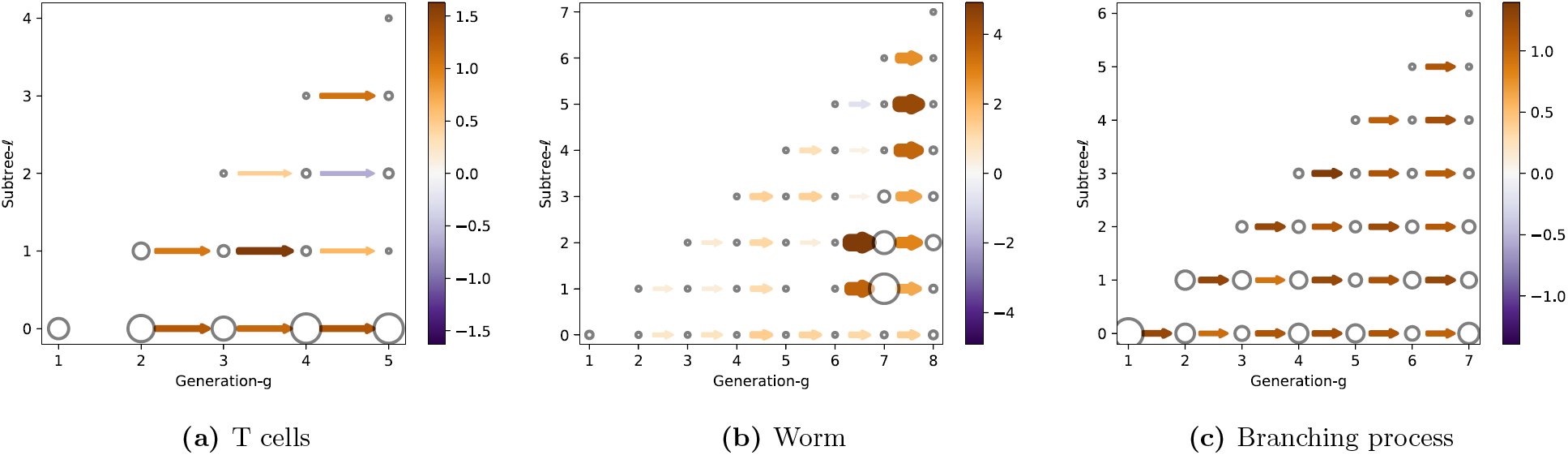
Dynamic lineage maps. These directed graphs in the natural variables show the dynamics of the bifurcated expression pattern in each subtree *𝓁*. The color (and thickness) of an edge between node *j* and *j*′ corresponds to the transmission strength, *β*_*jj*_′. The size of the node corresponds to the innovation strength, 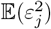.

As expected, the worm graph has the most structure. For example, transmission and innovation is small for the first few generations of each subtree, before “turning on” after generation 6. This means that the bifurcated expression of a subtree may be silent for many generations before appearing simultaneously over multiple descendants at a later generation. Note how transmission and innovations at *𝓁* = 0 are weak, illustrating how variation on the inter-pedigree level is small, as expected for a totipotent cell. Strong transmission is observed in particular subtrees at certain generations. For example, *β*_*jj*_′ is highest for *𝓁* = 2 between generations 6 and 7, and for *𝓁* = 5 between generations 7 and 8. We will discuss these features later when we assess the fate restriction associated with each subtree.

Although these characteristics could have been inferred just by viewing a single worm lineage directly, the point is that we now have a statistical method to extract such features from variable lineages. For example, the primary feature of the graph for T cells, which was not obvious from just looking at the lineages, is that subtree *𝓁* = 0 has the largest innovations and consistently strong transmission between generations (the exception is from generation 1, whose phenotype is not transmitted). This indicates that much of the variation is between pedigrees, rather than within the pedigree as it was for the worm. We will describe this in more detail in the next section.

Finally, we note that the graph for the branching process is featureless across all generations and in all subtrees, as would be expected for a stationary process.

### Fate profiles

Lineage variability maps describe the pattern of phenotypic associations throughout the lineage. However, as with lineage maps, our interest is often in using them to infer where fate is specified. In the introduction, we described how this involves identifying the most recent common ancestor of cells with shared fate. For a clonal pattern, where a cell fate is exclusive to a single subtree, we infer that fate was specified at (or near) a single lineal position - the root of that subtree. For a non-clonal pattern, which is likely for lineages with high variability, cell fate is expressed in multiple subtrees and we would infer that some fate was specified at multiple lineal positions. In *C. elegans* these inferences could be made visually [5]. Here we show how, by knowing the lineage variability map **∑**, we can make these inferences statistically, overcoming the problem of how to identify subtrees which shared phenotypes.

Before we begin, we must define what we mean by cell fate. In this study we define cell fate to be the measured phenotype of a cell at the latest generation studied, *G*. This practical definition allows us to analyze cell fate whether or not the phenotype in the last generation is actually a terminal fate. Also, by defining cell fate to be the phenotype itself rather than the cell type to which it is assigned, we can use the phenotypic measurements as is, without having to cluster or threshold them. Such discretization procedures can be difficult to define when phenotypes exist on a continuum of differentiation, as is often the case [77].

Having defined fate, we turn now to explaining its variability in terms of aspects of the lineage. We first partition the variability among the subtrees, or sources of variation. This quantifies how much of a cell’s fate is restricted by, or specified by, each subtree. We then examine the correlation of cell’s fate with the phenotypes of its ancestors. This identifies the generations over which a phenotypic fate has been stably expressed. Together these two measures, of fate restriction and fate expression, make up what we call fate profiles.

### Fate restriction by subtree

To determine how much cell fate is restricted by (i.e. specified by) each subtree, we partition the fate variability among the different sources of variation, each of which is located at the root of a *bifurcated* subtree. This is just the traditional problem of variance components analysis in nested groups (see Fig 5b). Since we have already calculated the spectral covariance matrix, we need only locate the appropriate components of variance along its diagonal (see Eq 16).

Consider the variance of a cell in generation *G*, given by *σ*_*GGG*_ (see Eq 5). This can be written as the the sum of independent contributions from each source (*𝓁, τ)*. These are known as the (normalized) components of variance in a classical ANOVA [78]. A convenient way to show this decomposition in our framework is to perform the inverse spectral transform of **∑**_Ω_ (for an example, see the Appendix A2.9). The result is given by

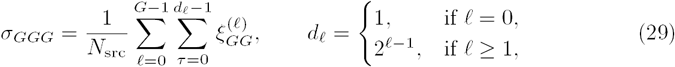

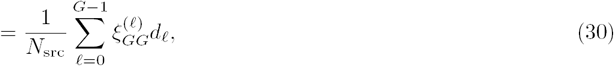

where *d*_*𝓁*_ is the number of transverse sources of variation at a given *𝓁*, and 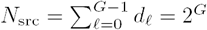 is the total number of sources of variation in a *G*-generation tree. The component of variance corresponding to source *𝓁* is thus given by 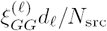 where 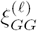 is found along the diagonal of **∑**_Ω_.

The resulting proportion of variance attributable to the *𝓁*-th source for a cell in generation *G* is given by

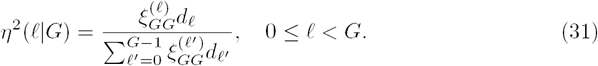

This measures the relative importance of each source of variation *𝓁* in explaining cell fate. Equivalently, it measures how much cell fate is restricted by subtree *𝓁.*

It will also be useful to calculate the cumulative proportion of total variance attributable to subtrees from 0 to *𝓁*, inclusive,

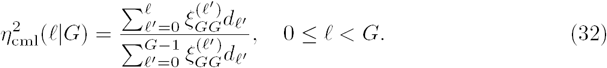

This gives a running total of the cell fate restricted by each successive subtree, starting at *𝓁* = 0 and is related to the intraclass correlation.

An obvious question is whether *η*^2^(*𝓁*| *G.*) would differ if we had simply performed a variance components analysis on the single generation *G*, ignoring measurements in the other generations. With complete data, our method would give the identical result to a variance components calculation: using a decomposable model for a Markov chain ensures that estimates of diagonal elements in **∑**_Ω_ (the components of variance) are given by the corresponding diagonal elements in ***S***_Ω_. If there were incomplete data however, data from other generations would help to estimate the missing data in generation *G*, improving the estimate of *η*^2^(*𝓁*|*G*).

### Fate expression by generation

Having determined how much fate is restricted by each subtree, we now determine how much cell fate is expressed by each generation. We do this by correlating the phenotype of a cell in generation *G* with those of its direct ancestors. The degree to which earlier generations are correlated with the last is a measure of when fate becomes expressed.

This definition of fate expression emphasizes the stability, or persistence, of a phenotypic fate rather than the absolute value of a phenotypic measurement. We have chosen this definition since our analysis should be general enough to work on data with substantial variability, where it may be difficult to define a cell fate in terms of some threshold level of expression.

Given a lineal position in generation *G* and its direct ancestor in generation *g*, the proportion of explained variance is just the squared correlation coefficient, or coefficient of determination,

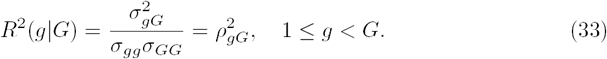

In the subscripts we have simplified the 3-index notation from Eq 5 by ignoring the third index. This does not cause confusion since in this context we are only concerned with direct ancestors.

Generalizing to prediction using multiple generations of direct ancestors up to and including that in generation *g* gives

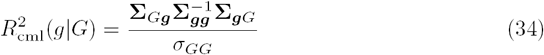

where ***g*** represents a vector of direct ancestors of the cell in generation *G* that are from generations 1 to *g* inclusive. Note that Eq 34 accounts for possible dependencies in the variation between ancestors. Unlike for the case of components of variance, contributions from different ancestral generations are not (in general) orthogonal.

### Comparing fate restriction and fate expression

Our measures of fate restriction and fate expression are complementary ways of explaining the variation of cell fate: *η*^2^(*𝓁*|*G*) explains fate in terms of shared ancestry (subtrees) while *R*^2^(*g* | *G*) explains fate in terms of ancestral phenotypes. We call these fate profiles. Both are plotted in Fig 11, with the top row giving the explained variance and the bottom row giving the cumulative explained variance.

**Fig 11.**
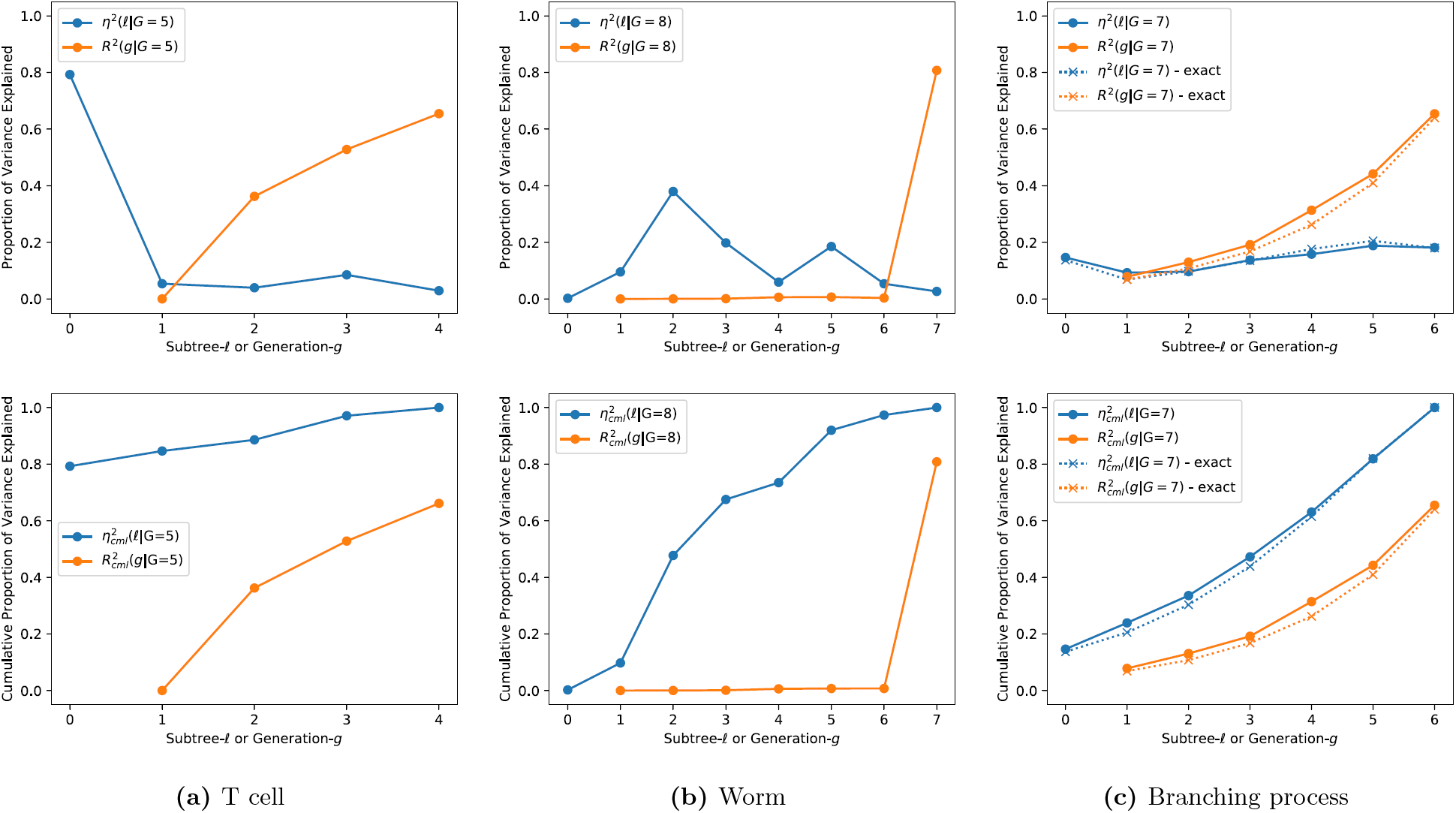
Fate profiles for different lineages. Explained variance (top row) and the cumulative explained variance (bottom row). *η*^2^(*𝓁* | *G*) (blue) measures how much the fate of a cell at generation *G* is restricted by each subtree *𝓁*. *R*^2^(*g* |*G*) (orange) measures how much a generation-*G* cell’s phenotype is correlated with its direct ancestor in generation *g*. Note that because of the Markov process is assumed to be first order (see Section ‘Sparsity’), 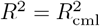 For the case of the simulated branching process the exact result is also shown to illustrate the accuracy of the inference procedure.

*η*^2^(*𝓁*|*G*) (blue line, top row) shows how much variation in *G* is restricted by each of the subtrees *𝓁*. For T cells, *𝓁* = 0 is by far the most important “subtree” for explaining fate (at *G* = 5). This is consistent with a cell that has limited potency, where the choice of founder cell severely restricts the range of fates available. In this case, any founder cell has already had 80% of its cell fate restricted. For the worm, cell fate is restricted by all subtrees *except 𝓁* = 0. Each zygote thus has 100% of its cell fate potential. This is consistent with the behavior for a totipotent cell. All subsequent subtrees contribute to cell fate, with *𝓁* = 2, 3, 5 being particularly important. This spread of fate specification over different subtrees might have been expected given the non-clonal expression pattern of *PHA-4*. While a clonal pattern is projected onto a single subtree, non-clonal patterns are projected onto multiple subtrees. For the branching process, contributions from all subtrees are comparable, as expected. Each subtree is, roughly speaking, equally important.

*R*^2^(*g*|*G*) (orange line, top row) gives the correlation of a cell in generation *G* with its direct ancestor in generation *g*. For T cells, *R*^2^ ≃ 0 for *g* = 1 indicating that, even though most cell fate (at least at *G* = 5) is set by the choice of founder cell, the founder does not actually resemble its descendants. For the worm, *R*^2^ ≃ 0 for 1 *≤ g ≤* 6. Thus none of the complicated structure in *η*^2^ for 0 *≤ 𝓁 ≤* 6 is reflected in *R*^2^.

This difference between fate restriction and fate expression is emphasized in the cumulative explained variance shown in the bottom row of Fig 11. For the worm, 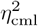 increases with each subtree (for *𝓁 >* 0) while 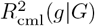 remains zero until *g* = 7. For cml cml the T cell, 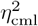 starts high at *𝓁*= 0, while 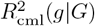 starts at zero and increases slowly with each generation. Contrast this with the branching process where 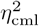 and 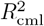 both start near zero and increase steadily in a similar fashion. Clearly a T cell lineage cannot be modeled as a branching process.

In the worm lineage, such *fate restriction before fate expression* captures what is perhaps obvious from the lineage map. Just by looking at Fig 1b we see how *PHA-4* expression is negligible until generation 7 whereupon it appears simultaneously across multiple subtrees. This implies that cells across those subtrees coordinated their fates before expressing them. Thus, for the worm, the fate profile merely restates, albeit in a quantitative way, what can be visualized in a single (invariant) pedigree. However, the advantage of the fate profile is that it can be applied to variable lineages, when simple visualization fails.

## Discussion

The lineage map, which has been instrumental in the discovery of fate specification mechanisms in simple organisms, was born from the study of invariant lineages and is not a particularly useful concept for understanding the more ubiquitous case of variable lineages. To address this, we have introduced lineage variability maps, which provide a way to describe lineages at the population level. Whereas the lineage map is a description of the pattern of phenotypes across a pedigree, the lineage variability map describes the pattern of phenotypic associations across a pedigree. This map of phenotypic associations, **∑**, provides quantitative answers to essential scientific questions such as those about cell potency, fate restriction, and the sources of variation in a lineage.

We have constructed lineage variability maps from a sample of highly-variable pedigrees from CD8^+^ T-lymphocytes up to five generations. These show that most of the variation in cell fate, defined here to be cell size at generation 5, is explained by the choice of naive cell. Yet, despite the pivotal role played by this founder in restricting cell fate, its phenotype is not predictive of fate: though a naive cell may specify that its descendants be large, it may not be large itself.

Although we expect to apply our technique primarily to variable pedigrees which are difficult to interpret by visualization alone, we can also apply it to invariant lineages to check our results. In fact, by constructing lineage variability maps from sample wild-type pedigrees from *C. elegans* marked for pharyngeal expression, we successfully recovered essential information in the known lineage map, identifying global features such as the small degree of inter-pedigree variation characteristic of a totipotent zygote, and the several-generation delay between fate specification and expression.

Yet our lineage variability maps capture important finer detail as well. Consider the peak in fate restriction at *𝓁* = 2 observed in Fig 11b. This arises from the strong bifurcation of fate traced back to the division of both P1 and of AB, progenitors located at *𝓁* = 2 (see Fig A1 for the labeling of lineal positions). That only a single daughter from P1 and from AB exhibit pharyngeal fate results in the spike in fate restriction that we observe. Interestingly, this phenomenon, of pharyngeal fate ensuing from two cousins at generation 3 (ABa and EMS) but not from their sisters, is a phenomenon that has been investigated in detail [79]. Such work laid the foundation for several further studies leading to a fundamental understanding of the molecular and cellular mechanisms for specification of pharyngeal tissue [80]. This demonstrates how, even though we may be ignorant of the ordering of the lineage, we can still detect a phenomenon of biological relevance that had previously required knowledge of this ordering. In other words, although we must assume lineage relationships are symmetric, this does not prevent us from detecting the effects of asymmetric lineage patterns from the ‘boost’ they give to the variance in particular subtrees.

Recent technological innovations have introduced a variety of methods for recording lineage data, involving both advanced imaging [19, 45–47] and genetic barcoding [20, 48, 50–56] techniques. With the statistical lineage mapping and fate profiling methods described in this manuscript, it should be possible to quantify several of the fundamental features of these lineages, such as the potency of progenitors, whether heterogeneity is clonal, and at what depth such heterogeneity appears. Just as the visual identification of fate bifurcations in the worm lineage map enabled the location of fate specification events to be discovered, the capacity to perform systematic screens to rapidly identify the important stages of fate restriction should contribute to a deeper understanding of the mechanisms of fate specification in more complex, more variable systems.

## Supporting information

**S1 Appendix Supplemental mathematical theory and derivations; nomenclature for *C. elegans* lineage.** Group symmetry and matrix decomposition

(A1), Group representation for a complete tree (A2), Maximum likelihood estimation (A3), and Lineage nomenclature for C. elegans (A4).

**S1 Data Lineage data.**

## Acknowledgments

We thank Alan Rubin for suggesting we test our method on the *C. elegans* lineage.

## Author contributions

**C**onceptualization DGH, SMR; **D**ata curation MY; **F**ormal analysis DGH, TPS; **F**unding acquisition DGH, SMR; **M**ethodology DGH, TPS; **R**esources MY; SMR; **S**oftware DGH; **S**upervision SMR; **W**riting - original draft preparation DGH; **W**riting - review and editing DGH, TPS, MY, SMR.

